# GSAlign – an efficient sequence alignment tool for intra-species genomes

**DOI:** 10.1101/782193

**Authors:** Hsin-Nan Lin, Wen-Lian Hsu

## Abstract

Personal genomics and comparative genomics are becoming more important in clinical practice and genome research. Both fields require sequence alignment to discover sequence conservation and variation. Though many methods have been developed, some are designed for small genome comparison while some are not efficient for large genome comparison. Moreover, most existing genome comparison tools have not been evaluated the correctness of sequence alignments systematically. A wrong sequence alignment would produce false sequence variants. In this study, we present GSAlign that handles large genome sequence alignment efficiently and identifies sequence variants from the alignment result. GSAlign is an efficient sequence alignment tool for intra-species genomes. It identifies sequence variations from the sequence alignments. We estimate performance by measuring the correctness of predicted sequence variations. The experiment results demonstrated that GSAlign is not only faster than most existing state-of-the-art methods, but also identifies sequence variants with high accuracy.

## Background

With the development of sequencing technology, the cost of whole genome sequencing is dropping rapidly. Sequencing the first human genome cost $2.7 billion in 2001; however, several commercial parties have claimed that the $1000 barrier for sequencing an entire human genome is broken [1]. Therefore, it is foreseeable that genome sequencing will become a reality in clinical practices in the near future, which brings the study of personal genomics and comparative genomics. Personal genomics involves the sequencing, analysis and interpretation of the genome of an individual. It can offer many clinical applications, particularly in the diagnosis of genetic deficiencies and human diseases [2]. Comparative genomics is another field to study the genomic features of different organisms. It aims to understand the structure and function of genomes by identifying regions with similar sequences between characterized organisms.

Both personal genomics and comparative genomics require sequence alignment to discover sequence conservation and variation. Sequence conservation patterns can be helpful to predict functional categories, whereas variation can be helpful to infer relationship between organisms or populations in different areas. Studies have shown that variation is important to human health and common genetic disease [3–5]. The alignment speed is an important issue since a genome sequence usually consists of millions of nucleotides or more. Methods based on the traditional alignment algorithms, like AVID [6], BLAST [7] and FASTA [8], are not able to handle large scale sequence alignment. Many genome comparison algorithms have been developed, including ATGC [9, 10], BBBWT [11], BLAT [12], BLASTZ [13], Cgaln [14], chainCleaner [15], Harvest [16], LAST [17], MAGIC [18], MUMmer [19–22], and minimap2 [23].

One of important applications of genome comparison is to identify sequence variations between genomes, which can be found by linearly scanning their alignment result. However, none of the above-mentioned methods have been evaluated the correctness of sequence alignment regarding variation detection. A wrong sequence alignment would produce false sequence variants. In this study, we estimated the performance of each selected genome sequence comparison tool by measuring the correctness of sequence variation. Below we briefly describe the algorithm behind each method.

AVID finds maximal matches between two sequences using a suffix tree structure. It then clusters all the matches and split the sequences accordingly. Each cluster forms a local alignment of the two input sequences. BBBWT is the first attempt to adopt Burrows Wheeler Transformation (BWT) [24] on genomic sequence search. However, BBBWT is only designed to find all bi-unique *k*-mers in common between two genomes. It does not produce any sequence alignments. BLAT adopts a BLAST-like algorithm that rapidly scans for relatively short matches and extends those into high-scoring pairs. BLASTZ modifies the algorithm of Gapped BLAST to align long genomic sequences. It first finds short near-exact matches, extends each short match without allowing gaps, and then extends these again by a dynamic programming procedure that permits gaps. LASTZ is a drop-in replacement for BLASTZ. It adopts similar alignment procedure, though it includes more seeding strategies and reduces memory requirements.

Cgaln divides the input sequences into blocks with a fixed length. It then performs block-to-block similarity evaluation that is based on the frequency of common seeds in the blocks. The nucleotide-level alignment is derived from the block-level alignments with a seed-extension strategy. The chainCleaner aims to compare orthologous genes between the human and other vertebrate genomes. Harvest also focuses on core-genome alignment for microbial genomes. The alignment is based on a suffix tree to identify maximal unique matches (MUMs). It then uses MUMs to recruit similar genomes and anchor the multiple alignment. LAST follows the seed-and-extend approach, but it uses adaptive seeds to increase both alignment sensitivity and speed for large sequence comparison. Adaptive seeds are matches that are chosen based on their rareness. MUMmer was the first aligner to use suffix trees to find potential anchors for genome sequence alignments. One of the two input genome sequences is used as the reference sequence to build the suffix tree, and the second one is used to query against the tree. MUMmer identifies all maximal matches between the two sequences and then finds the longest increasing subsequence (LIS) from the sorted matches. The pairwise sequence alignment is built on the LIS by performing the Smith-Waterman alignment to close gaps between the ordered matches. MUMmer was shown to be 4 to 110 times faster than AVID, BLASTZ, and LAGAN [25]. Therefore, MUMmer has become a popular algorithm for comparing genome sequences in recent years. Minimap2 is a multi-purpose alignment tool. It follows a “seed-chain-align” procedure. It collects minimizers of the reference sequences and creates a hash table to index those minimizers. Then Mimimap2 finds all query minimizers to find exact matches to the reference, and identifies co-linear anchors as chain. Finally, Mimimap2 applies dynamic programming to extend from the ends of chains and to close gaps between adjacent anchors in chains.

Recently, many NGS read mapping algorithms use BWT or FM-index [26] to build an index for the reference sequences and identify maximal exact matches by searching against the index array with a query sequence. It has been shown that BWT-based read mappers are more memory efficient than hash table based mappers [27]. In this study, we used BWT to perform seed exploration for genome sequence alignment. We demonstrated that GSAlign is efficient in finding both exact matches and differences between two intra-species genomes. The differences include all single nucleotide polymorphisms (SNPs), insertions, and deletions. Moreover, the alignment is ultra-fast and memory efficient. The source code of GSAlign is available at https://github.com/hsinnan75/GSAlign.

## Methods

The algorithm of GSAlign is derived from our DNA read mapper, Kart [28]. Kart adopts a divide-and-conquer strategy to separate a read into regions with and without differences. The same strategy is applicable to genome sequence alignment. However, in contrast with NGS short read alignment, genome sequence alignment often consists of multiple subalignments that are separated by dissimilar regions or variants. In this study, we present GSAlign for handling genome sequence alignment.

### Algorithm overview

Similar to most existing methods, GSAlign also follows the “seed-chain-align” procedure to perform genome sequence alignment. However, the details of each step are quite different. GSAlign consists of three main steps: *LMEM identification (seed)*, *similar region identification (chain)*, and *alignment processing (align)*. We define a *local maximal exact match* (LMEM) as a common substring between two genomes that begins at a specific position of query sequence. In the LMEM identification step, GSAlign finds LMEMs with variable lengths and then converts those LMEMs into simple pairs. A simple pair represent two identical sequence fragments, one from the reference and one from the query sequence. In the similar region identification, GSAlign clusters those simple pairs into disjoint groups. Each group represents a similar region. GSAlign finds all local gaps in each simple region. A local gap (defined as a normal pair) is the gap between two adjacent simple pairs. In the alignment-processing step, GSAlign closes local gaps to build a complete local alignment for each similar region and identifies all sequence variations during the process. Finally, GSAlign outputs the alignments of all similar regions, a VCF (variant call format) file, and a dot-plot representation (optional). The contribution of this study is that we optimize those steps and integrate them into a very efficient algorithm that saves both time and memory and produces reliable alignments.

### Burrows-Wheeler transform

We give a brief background of BWT algorithm below. Consider a text *T* of length *L* over an alphabet set Σ; *T* is attached with symbol *$* at the end, and *$* is lexicographically smaller than any character in Σ. Let *SA*[0, *L*] be the suffix array of *T*, such that *SA*[*i*] indicates the starting position of the *i*-th lexicographically smallest suffix. The BWT of *T* is a permutation of *T* such that *BWT*[*i*] = *T*[SA[*i*] − 1] (Note that if *SA*[*i*] = 0, *BWT*[*i*] = *$*). Given a pattern *S*, suppose SA[*i*] and SA[*j*] are the smallest and largest suffices of *T* where *P* is their common prefix, the range [*i*, *j*] indicates the occurrences of *S*. Thus, given an SA range [*i*, *j*] of pattern *P*, we can apply the backward search algorithm to find the SA range [*p*, *q*] of *zP* for any character *z*. If we build the BWT with the reverse of *T*, the backward search algorithm can be used to test whether a pattern *P* is an exact substring of *T* in *O*(|*P*|) time by iteratively matching each character in *P*. One of the BWT index algorithms was implemented in BWT-SW [29] and it was then modified to work with BWA [27]. For the details of BWT index algorithm and the search algorithm, please refer to the above-mentioned methods and Kart.

### LMEM identification

Given two genome sequences *P* and *Q*, GSAlign generates the BWT array with *P* and its reverse complementary sequence *P’*. Let *P*[*i*_1_] be the *i*_1_-th nucleobase of *P*, and *P*[*i*_1_, *i*_2_] be the sequence fragment between *P*[*i*_1_] and *P*[*i*_2_]. GSAlign finds LMEMs by searching against the BWT array with *Q*. Since each LMEM is a common substring that begins at a specific position of *Q*, it is represented as a simple region pairs (abbreviated as *simple pairs*) in this study and denoted by a 4-tuple (*i*_1_, *i*_2_, *j*_1_, *j*_2_), meaning *P*[*i*_1_, *i*_2_] = *Q*[*j*_1_, *j*_2_] and *P*[*i*_2+1_] ≠ *Q*[*j*_2+1_]. If the common substring appear multiple times (i.e., frequency > 1), it would be transformed into multiple simple pairs. For example, if the substring *Q*[*j*_1_, *j*_2_] is identical to *P*[*i*_1_, *i*_2_] and *P*[*i*_3_, *i*_4_], it would be represented as two simple pairs (*i*_1_, *i*_2_, *j*_1_, *j*_2_) and (*i*_3_, *i*_4_, *j*_1_, *j*_2_). Note that an LMEM is transformed into simple pairs only if its size is not smaller than a user-defined threshold *k* and its occurrences are less than *f*. We investigate the effect of threshold *k* and *f* in the Supplementary data and we found that GSAlign performs equally well with different thresholds.

The BWT search iteratively matches every nucleotide of the query genome *Q*. It begins with *Q*[*j*_1_] (*j*_1_ = 0 at the first iteration) and stops at *Q*[*j*_2_] if it meets a mismatch at *Q*[*j*_2+1_], i.e., the SA range of *Q*[*j*_1_, *j*_2+1_] = 0. The next iteration of BWT search will start from *Q*[*j*_2_+1] until it meets another mismatch.

Fig. 1 illustrates the simple pairs between *P* and *Q*. Each rectangle is a simple pair and the width is the size of the simple pair. Note that the LMEM identification can be processed simultaneously if GSAlign runs using multiple threads. For each query sequence in *Q*, if there are *N* threads, GSAlign divides it into *N* blocks of equal size and each thread identifies LMEMs for a sequence block independently. The simple pairs identified by each thread will be merged together afterward. The multithreading can be also applied in the following alignment step. We will demonstrate that such parallel processing greatly speedup the alignment process.

**Fig. 1.**
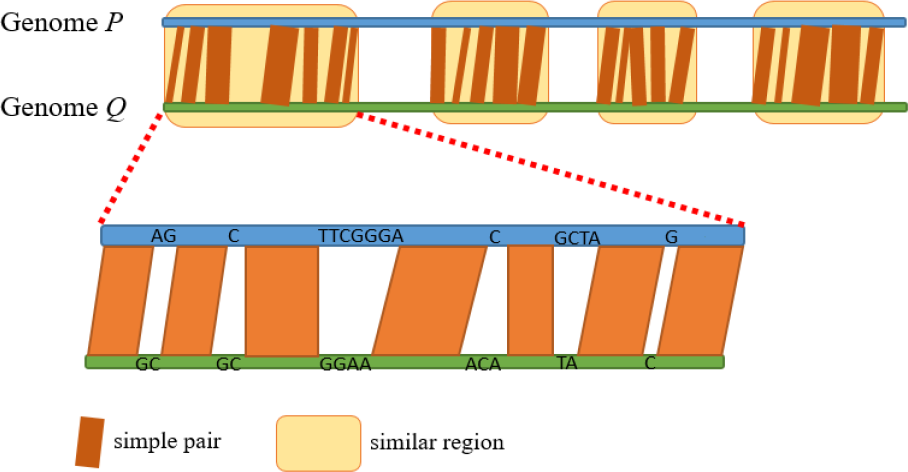
The simple pairs between genome P and Q. Each rectangle is a simple pair and the width is the size of the simple pair. A similar region is a group of neighboring simple pairs.

### Similar region identification

After collecting all simple pairs in the previous step, GSAlign sorts the simple pairs according to their position differences between genomes *P* and *Q* and clusters those into disjoint groups. The clustering algorithm is described below.

Suppose *S*_*k*_ is a simple pair (*i*_*k*,1_, *i*_*k*,2_, *j*_*k*,1_, *j*_*k*,2_), we define *PosDiff*_*k*_ = *i*_*k*,1_ − *j*_*k*,1_. If two simple pairs have similar *PosDiff*, they are co-linear. We sort all simple pairs by their *PosDiff* to group all co-linear simple pairs. The clustering starts with the first simple pair *S*_1_ and we check if the next simple pair (*S*_2_) is within a threshold *MaxDiff* (the default value is 25). The size of *MaxDiff* determines the maximum indel size allowed between two simple pairs. If |*PosDiff*_*1*_ − *PosDiff*_*2*_| ≤ *MaxDiff*, we then check the PosDiff of *S*_*2*_ and *S*_*3*_ until we find two simple pairs *S*_*k*_ and *S*_*k*+*1*_ whose |*PosDiff*_*k*_ − *PosDiff*_*k*+*1*_| > *MaxDiff*. In such case, the clustering breaks at *S*_*k*+*1*_ and simple pairs *S*_*1*_, *S*_*2*_, …, *S*_*k*_ are clustered in the same group. We then sort *S*_*1*_, *S*_*2*_, …, *S*_*k*_ by their positions at sequence Q (i.e., the third value of 4-tuple). Since simple pairs are re-sorted by their positions at sequence Q, some of them may be not co-linear with their neighboring simple pairs and they are considered as outliers. We remove those outliers from the simple pair group. For those simple pairs of same positions at sequence Q, we keep the one with the minimal difference of *PosDiff* compared to the closest unique simple pair. Then we check every two adjacent simple pairs *s*_*a*_ = (*i*_a,1_, *i*_a,2_, *j*_a,1_, *j*_a,2_) and *s*_*b*_ = (*i*_b,1_, *i*_b,2_, *j*_b,1_, *j*_b,2_), we define gap(*S*_*a*_, *S*_*b*_) = *j*_*b*,1_ − *j*_*a*,2_. If gap(*S*_*a*_, *S*_*b*_) is more than 300bp and the sequence fragments in the gap are dissimilar, we consider *S*_*b*_ as a break point of a simple region. We investigate different gap size thresholds in the Supplementary data and found that GSAlign was not sensitive to the threshold. To determine whether the sequence fragment in a gap are similar, we use k-mers to estimate their similarity. If the number of common k-mers is less than gap(*S*_*a*_, *S*_*b*_) / 3, they are considered dissimilar. In such case, we consider *S*_*b*_ as a break point of a simple region. GSAlign then continues the clustering with *S*_*k*+*1*_ and identifies more similar regions for the remaining simple pairs until all simple pairs are visited.

We use a toy example to illustrate the process of simple pair clustering and outlier removing. Suppose GSAlign identifies nine simple pairs as shown in Fig 2(A). We sort these simple pairs by their PosDiff and start clustering process with S_1_. Simple pairs *S*_*1*_, *S*_*2*_, …, *S*_*8*_ are clustered in the same group since any two adjacent simple pairs in the group have similar PosDiff. For example, |*PosDiff*_*1*_ − *PosDiff*_*2*_| = 10, and |*PosDiff*_*2*_ − *PosDiff*_*3*_| = 0. By contrast, |*PosDiff*_*8*_ − *PosDiff*_*9*_| = 60, therefore we break the grouping at *S*_*9*_. We then re-sort *S*_*1*_, *S*_*2*_, …, *S*_*8*_ by their positions at sequence Q as shown in Fig 2(B), and mark S_6_ and S_7_ are not unique since the two simple pairs are from the same position at Q. We remove S_1_ and S_8_ since they are not co-linear with their neighboring simple pairs. Then we compare S_6_ and S_7_ and keep S_6_ because it has the minimal difference of *PosDiff* with its neighboring unique simple pairs. Finally, we confirm there is no any large gap between any two adjacent simple pairs in the group. Thus, S_3_, S_1_, S_6_, S_2_, S_4_, S_5_, and S_8_ forms a simple region, and upon which we can generate a local alignment.

**Fig. 2.**
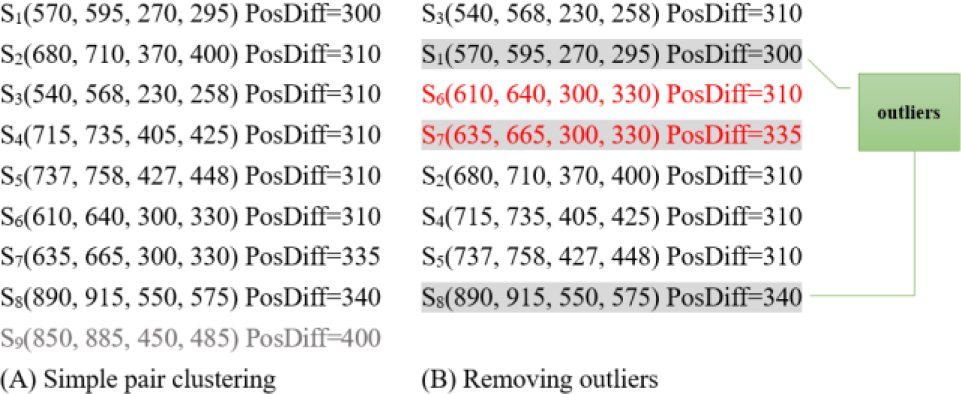
A toy example illustrating the process of simple clustering and outlier removing.. GSAlign clusters simple pairs and remove outliers according to PosDiff. Simple pairs in red are not unique. Simple pairs with gray backgrounds are considered as outliers and they are removed from the cluster.

Given two adjacent simple pairs in the same cluster, *s*_*a*_ = (*i*_a,1_, *i*_a,2_, *j*_a,1_, *j*_a,2_) and *s*_*b*_ = (*i*_b,1_, *i*_b,2_, *j*_b,1_, *j*_b,2_), we say *s*_*a*_ and *s*_*b*_ overlap if *i*_a,1_ ≤ *i*_b,1_ ≤ *i*_a,2_ or *j*_a,1_ ≤ *j*_b,1_ ≤ *j*_a,2_. In such cases, the overlapping fragment is chopped off from the smaller simple pair. For example, BWT index. Fig 3. shows a tandem repeat with different copies in genome *P* and *Q*. In this example, “ACGT” is a tandem repeat where *P* has seven copies and *Q* has nine copies. GSAlign identifies two simple pairs in this region: *A*(301, 330, 321, 350) and *B*(323, 335, 351, 363). *A* and *B* overlap between *P*[323, 330]. In such cases, we remove the overlap from the preceding simple pair (i.e., *A*). After removing the overlap, *A* becomes (301, 322, 321, 342) and we create a gap of *Q*[343, 350]. After removing overlaps, we check if there is a gap between any two adjacent simple pairs in each similar region. We fill gaps by inserting normal pairs. A normal pair is also denoted as a 4-tuple (*i*_1_, *i*_2_, *j*_1_, *j*_2_) in which *P*[*i*_1_, *i*_2_] ≠ *Q*[*j*_1_, *j*_2_] and the size of *P*[*i*_1_, *i*_2_] or *Q*[*j*_1_, *j*_2_] can be 0 if one of them is an deletion. Suppose we are given two adjacent simple pairs (*i*_2*q*−1_, *i*_2*q*_, *j*_2*q*−1_, *j*_2*q*_) and (*i*_2*q*+1_, *i*_2*q*+2_, *j*_2*q*+1_, *j*_2*q*+2_). If *i*_2*q*+1_ − *i*_2*q*_ > 1 or *j*_2*q*+1_ − *j*_2*q*_ > 1, then we insert a normal pair (*i*_*r*_, *i*_*r*+1_, *j*_*r*_, *j*_*r*+1_) to fill the gap, where *i*_*r*_ − *i*_2*q*_= *i*_2*q*+1_ − *i*_*r*+1_ = 1 if *i*_2*q*+1_ − *i*_2*q*_ > 1; otherwise *i*_*r*_ = *i*_*r*+1_ = −1 meaning the corresponding fragment size is 0. Likewise, *j*_*r*_ − *j*_2*q*_ = *j*_2*q*+1_ − *j*_*r*+1_ = 1 if *j*_2*q*+1_ − *j*_2*q*_ > 1, otherwise let *j*_*r*_ = *j*_*r*+1_ = −1.

**Fig. 3.**
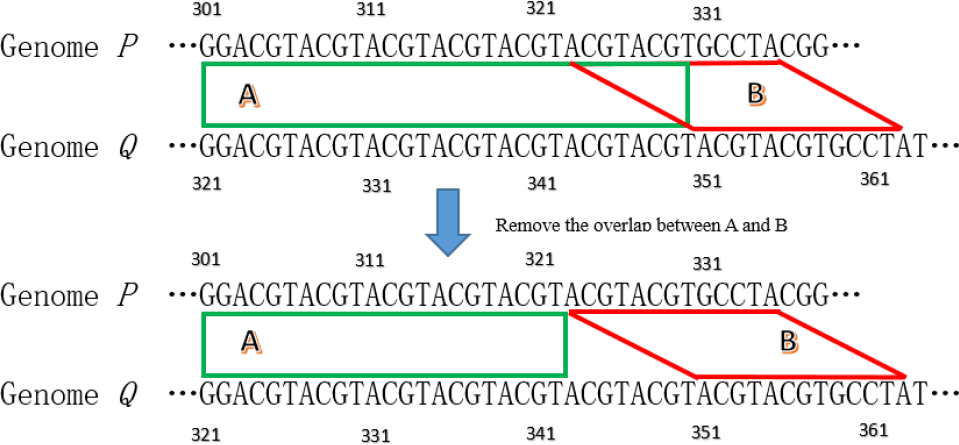
Simple pairs A and B overlaps due to tandem repeats of “ACGT”. We remove the overlapped fragment from simple pair A (the preceding one).

### Alignment processing

At this point, GSAlign has identified similar regions that consist of simple pairs and normal pairs. In this step, GSAlign only focuses on normal pairs. If the sequence fragments in a normal pair have equal size, it is very likely the sequence fragments only contains substitutions and the un-gapped alignment is already the best alignment; if the sequence fragments contain indels, gapped alignment is required. Therefore, we classify normal pairs into the following types:

1. A normal pair is *Type I* if the fragment pair has equal size and the number of mismatches in a linear scan is less than a threshold;
2. A normal pair is *Type II* if one of the fragment is a null string and the other contains at least one nucleobase;
3. The remaining normal pairs are *Type III;*

Thus, only *Type III* require gapped alignment. GSAlign applies the KSW2 algorithm [30] to perform gapped alignment. The alignment of each normal pair is constrained by the sequence fragment pair. This allows GSAlign to generate their alignments simultaneously with multiple threads. At the end, the complete alignment of the genome sequences is the concatenation of the alignment of each simple and normal pairs.

### Differences among GSAlign, MUMmer4, and Minimap2

In general, GSAlign, MUMmer4, and Minimap2 follow the conventional seed-chain-align procedure to align genome sequences. However, the implementation details are very different from each other. MUMmer4 combines the ideas of suffix arrays, the longest increasing subsequence (LIS) and Smith-Waterman alignment. Minimap2 uses minimizers (k-mers) as seeds and identifies co-linear seeds as chains. It applies a heuristic algorithm to cluster seeds into chains and it uses dynamic programming to closes between adjacent seeds. GSAlign integrates the ideas of BWT arrays, PosDiff-based clustering and dynamic programming algorithm. GSAlign divides the query sequence into multiple blocks and identifies LMEMs on each block simultaneously using multiple threads. More importantly, GSAlign classifies normal pairs into three types and only *Type III* normal pairs require gapped alignment. This divide-and-conquer strategy not only reduces the number of fragment pairs requiring gapped alignment, but also shortens gap alignment sizes. Furthermore, GSAlign can produce the alignments of normal pairs simultaneously with multi-threads. Though MUMmer4 supports multi-threads to align query sequences in parallel, the concurrency is restricted to the number of sequences in the query.

## Results

### Experiment design

GSAlign takes two genome sequences: one is the reference genome for building the BWT index, and the other is the query genome for searching against the BWT array. If the reference genome has been indexed before-hand, GSAlign can read the index directly. After comparing the genome sequences, GSAlign outputs all local alignments in MAF format or BLAST-like format, a VCF file, and a dot-plot representation for each query sequence.

The correctness of sequence alignment is an important issue and variant detection is one of the major applications for genome sequence alignment. Therefore, we estimate the correctness of sequence alignments by measuring the variant detection accuracy. Though most of genome alignment tools do not output variants, we can find variants out by linearly scanning the sequence alignments. This novel measurement is sensitive to misalignments, thus we consider it is a fair measurement to estimate the performance of sequence alignment.

We randomly generates sequence variations with the occurrences of 20,000 substitutions (SNVs), 350 small indels (1~10 bp), 100 large indels (11~20 bp) for every 1M base pairs. To increase the genetic distance, we generate different frequencies of SNVs. Benchmark datasets labelled with 1X contain around 20,000 SNVs for every 1M base pairs, whereas datasets labelled with 3X (or 5X) contain 60,000 (or 100,000) SNVs per million bases. We generate three synthetic datasets with different SNV frequencies using the human genome (GRCh38). The synthetic datasets are referred to as simHG-1X, simHG-3X, and simHG-5X, respectively. To evaluate the performance of genome sequence alignment on real genomes, we download the diploid sequence of NA12878 genome and the variant calls (the sources are shown in Supplementary data). The diploid sequence of NA12878 consists of 3,088,156 single nucleotide variants (SNVs) and 531,315 indels derived from NGS data analysis. The reference variants are generated from NGS data analysis. Please note that GSAlign is a genome alignment tool, rather than a variant caller such as Freebayes or GATK HaplotypeCaller. GSAlign identifies sequence variants from genome sequence alignment, while Freebayes and GATK HaplotypeCaller identify variants from NGS short read alignments. We use sequence variants to estimate the correctness of sequence alignment in this study. Table 1 shows the genome size and their variation numbers of each benchmark dataset.

**Table 1.** The synthetic datasets and the number of simulated sequence variations.

| Dataset | Genome size | SNV | small indel | large indel | Total variants |
| --- | --- | --- | --- | --- | --- |
| simHG-1X | 3,088,279,342 | 58,421,383 | 1,001,626 | 285,757 | 59,708,766 |
| simHG-3X | 3,088,292,247 | 175,100,939 | 962,721 | 275,584 | 176,339,244 |
| simHG-5X | 3,088,289,999 | 291,714,646 | 919,762 | 263,271 | 292,897,679 |
| NA12878 | 6,070,700,436 | 3,088,156 | 531,315 | NA | 3,619,471 |

In this study, we compare the performance of GSAlign with several existing genome sequence aligners, including LAST (version 828), Minimap2 (2.17-r943-dirty), and MUMmer4 (version 4.0.0beta2). We exclude the others because they are either unavailable or developed for multiple sequence alignments, like Cactus [31], Mugsy [32], or MULTIZ [33]. We exclude BLAT because it fails to produce alignments for larger sequence comparison; we exclude LASTZ because it does not support multi-thread. Moreover, LASTZ fails to handle human genome alignment.

### Measurement

We define a true positive case (TP) as a true sequence variation identified from sequence alignment; a false positive case (FP) as a false sequence variation; and a false negative case (FN) as a true sequence variation that is not identified. A predicted SNV event is considered true if the genomic coordinate is exactly identical to the true event; a predicted indel event is considered true if the predicted coordinate is within 10 nucleobases of the true event.

To estimate the performance for existing methods, we filter out sequence alignments whose sequence identity are lower than a threshold (for Mummer4 and LAST) or quality score are 0 (for Minimap2). The argument setting used for each method are shown in the Supplementary. We estimate the precision and recall on the identification of sequence variations for each dataset. GSAlign, Minimap2, MUMmer4, and LAST can load premade reference indexes; therefore, we run these methods by feeding the premade reference indexes and they are running with 8 threads.

### Performance evaluation on synthetic datasets

Table 2 summarizes the performance result on the three synthetic datasets. It is observed that GSAlign and Minimap2 have comparable performance on the benchmark dataset. Both produce alignments that indicate sequence variations correctly. MUMmer4 and LAST produce less reliable alignments than GSAlign and Minimap2. Though we have filtered out some of alignments based on sequence identity, their precisions and recalls are not as good as those of GSAlign and Minimap2. In particular, the precision of indel events of MUMmer4 and LAST are much lower on the dataset of simHG-5X. It implies that the two methods are not designed for genome sequence alignments with less sequence similarity. We also compare the total number of local alignments each method produces for the benchmark datasets. It is observed that GSAlign produces the least number of local alignments, though it still covers most of the sequence variants. For example, GSAlign produces 250 local alignments for simHG-1X, whereas the other three methods produce 417, 3111 and 1168 local alignments, respectively.

**Table 2.** The performance evaluation on the three HG38 synthetic data sets.

| Dataset | Method | SNV |  | Indel |  | Local | Run |
| --- | --- | --- | --- | --- | --- | --- | --- |
|  |  | precision | recall | precision | recall | align# | time (min) |
| SimHG-1X | GSAAlign | 1.000 | 1.000 | 0.999 | 0.999 | 250 | 11 |
|  | Minimap2 | 1.000 | 0.996 | 0.999 | 0.995 | 417 | 39 |
|  | MUMmer4 | 0.998 | 0.932 | 0.985 | 0.932 | 3111 | 869 |
|  | LAST | 1.000 | 0.992 | 0.992 | 0.947 | 1168 | 2524 |
| SimHG-3X | GSAAlign | 1.000 | 0.998 | 0.994 | 0.997 | 366 | 18 |
|  | Minimap2 | 1.000 | 0.996 | 0.991 | 0.995 | 561 | 37 |
|  | MUMmer4 | 0.989 | 0.923 | 0.796 | 0.925 | 4925 | 289 |
|  | LAST | 1.000 | 0.990 | 0.809 | 0.950 | 1234 | 1185 |
| SimHG-5X | GSAAlign | 1.000 | 0.993 | 0.958 | 0.992 | 587 | 24 |
|  | Minimap2 | 1.000 | 0.995 | 0.952 | 0.994 | 1058 | 40 |
|  | MUMmer4 | 0.986 | 0.907 | 0.486 | 0.912 | 5513 | 157 |
|  | LAST | 1.000 | 0.981 | 0.461 | 0.947 | 1636 | 458 |

In terms of runtime, it can be observed that GSAlign spends the least amount of runtime on the three datasets. Minimap2 is the second fastest method. Though MUMmer4 is faster than LAST, it produces worse performance than LAST. We observe that LAST is not very efficient with multi-threading. Though it runs with eight threads, it only uses single thread most of the time during the sequence comparison. Interestingly, GSAlign spends more time on less similar genome sequences (ex. simHG-5X) because there are more gapped alignments, whereas MUMmer4 and LAST spends more time on more similar genome sequences (ex. simHG-1X) because they handle more number of seeds. Minimap2 spends similar amount of time on the three synthetic datasets because Minimap2 produces similar number of seeds for those datasets. Note that it is possible to speed up the alignment procedure by optimizing the parameter settings for each method; however it may complicate the comparison.

### Performance evaluation on NA12878

The two sets of diploid sequence of NA12878 are aligned separately and the resulting VCF files are merged together for performance evaluation. Because many indel events of NA12878 locate in tandem repeat regions, we consider a predicted indel is a TP if it locates at either end of the repeat region. For example, the two following alignments produce identical alignment scores:

It can be observed that the two alignments produce different indel events.

~~~
AGCATGCATTG AGCATGCATTG
AGCAT----TG, and AG----CATTG.
~~~

In such case, both indel events are considered true positives if one of them is a true indel. Table 3 summaries the performance evaluation on the real dataset. It is observed that GSAlign, Minimap2 and LAST produce comparable results on SNV and indel detection. They have similar precisions and recalls. MUMmer4 produces worse performance on SNV and indel detection. Its precision and recalls are not as good as the other three methods’. In terms of runtime, it is observed that GSAlign only spends 5 minutes to align the diploid sequences of NA12878 with HG38. Minimap2 is the second fastest method. It spends 65 minutes. LAST and MUMmer4 spend 1305 and 3898 minutes, respectively. In terms of memory consumption, it is observed that GSAlign consumes the least amount of memory among the selected methods. It requires 14 GB to perform the genome comparison, while MUMmer4 requires 57 GB.

**Table 3.** The performance evaluation on HG38 and the diploid sequence of NA12878.

| Dataset | Method | SNV |  | Indel |  | Run | Memory |
| --- | --- | --- | --- | --- | --- | --- | --- |
|  |  | Precision | Recall | Precision | Recall | Time (min) | usage (GB) |
| NA12878 (Diploid) | GSAAlign | 0.832 | 0.969 | 0.759 | 0.767 | 5 | 14 |
|  | Minimap2 | 0.830 | 0.970 | 0.754 | 0.768 | 65 | 23 |
|  | MUMmer4 | 0.752 | 0.946 | 0.711 | 0.749 | 3898 | 57 |
|  | LAST | 0.832 | 0.969 | 0.760 | 0.764 | 1305 | 28 |

### Sequence comparison between difference species

Though GSAlign is designed for comparing intra-species genomes, it can be used to identify conserved syntenic regions for inter-species genomes. Here we compare human genomes with whole chimpanzee genome and mouse chromosome 12. We compare human (HG38) and chimpanzee (PanTro4) genomes using GSAlign with 8 threads. It spends around nine minutes on performing the genome comparison. GSAlign generates 98298 local alignments whose total length is 2,253M bases (PanTro4 contains 390M undetermined bases ‘N’). GSAlign identifies 29.2 million SNVs and 3.4 million indels between HG38 and panTro4.

Mouse chromosomes share common ancestry with human chromosomes [34]. Here we demonstrate the sequence comparison between human genome and mouse chromosome 12 by showing the dot-plot matrix generated by GSAlign. Though the genome sequences of the two species are very dissimilar, they still share conservation of genetic linkage groups. In this analysis, GSAlign spends three minutes to compare HG38 and mouse chromosome 12 and it generates 2,713 local alignments with a total length of 1,738K bases. Among all the 22 human chromosomes, GSAlign discovers that human chromosomes 2, 7 and 14 share the largest number of conserved syntenic segments with mouse chromosome 12. GSAlign visualizes their similarity with a dot-plot presentation. Fig. 4 shows the dot-plot of those chromosomes. Comparing the result with existing studies, we find that the dot-plot is consistent with Fig. 4(f) in the study of Mouse Genome Sequencing Consortium [34].

**Fig. 4.**
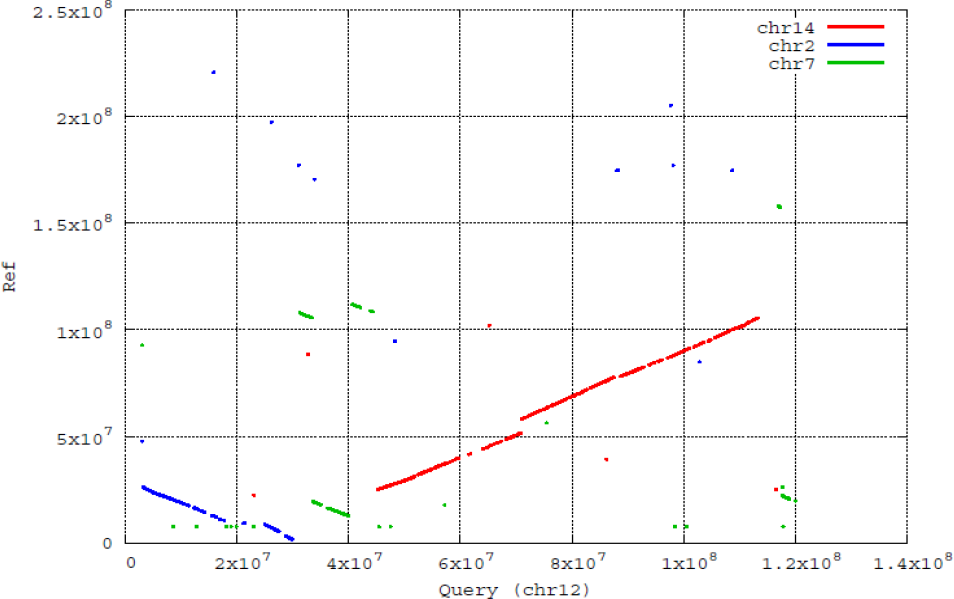
The dotplot of the alignment for human chromosomes 2, 7, and 14 and mouse chromosome 12. Each line or dot indicates a local alignment.

## Conclusions

In this study, we propose GSAlign to handle genome sequence comparison and evaluate the correctness of sequence alignment by measuring the accuracy of variant detection. We adopt the divide-and-conquer strategy to divide the genome sequences into gap-free fragment pairs and gapped fragment pairs. GSAlign is a BWT-based genome sequence aligner. Therefore, it requires less amount of memory than hash table-based or tree-based aligners do. GSAlign also supports parallel computing for genome sequence comparison, thus it is more efficient when comparing large genomes. We evaluate the performances of GSAlign with synthetic and real datasets. The experiment result shows that GSAlign is the fastest among the selected methods and it produces perfect or nearly perfect precisions and recalls on the identification of sequence variations for most of the datasets.

With the emergence of personal genomics and comparative genomics, we believe GSAlign can be a useful tool. It shows the abilities of ultra-fast alignment as well as high accuracy and sensitivity for detecting sequence variations. As more genome sequences become available, the demand for genome comparison is increasing. Therefore an efficient and robust algorithm is most desirable.

## Acknowledgements

Funding: This work was supported by the Bioinformatics Core Facility for Translational Medicine and Biotechnology Development from the Ministry of Science and Technology (MOST) of Taiwan under grant 105-2319-B-400-002.

## Conflict of Interest

none declared.

