## Supplementary data for "GSAlign – an efficient sequence alignment tool for intra-species genomes"

Hsin-Nan Lin and Wen-Lian Hsu
Institute of Information Science, Academia Sinica, Taipei, Taiwan

### Supplementary Data

#### 1. The effect of MaxPosDiff threshold

The size of MaxPosDiff determines the maximum indel size is allowed between adjacent seeds (simple pairs). We separate two adjacent simple pairs $S_a$ and $S_b$ into separate clusters if $|PosDiff_a - PosDiff_b| \geq MaxPosDiff$. The default value of MaxPosDiff is 25. Here, we investigate the effect of MaxPosDiff. Table *S1* summaries the analysis result. We tested MaxPosDiff between 25 and 50 on Sim_Chr1. It can be observed that GSAlign performs equally well with different thresholds.

*Table S1*. The effect of *MaxPosDiff* for GSAlign on the Sim_Chr1 dataset.

| ***Sim_Chr1*** | ***MaxPosDiff*** | ***SNV*** | | ***Indel*** | | ***Run time (second)*** |
|---|---|---|---|---|---|---|
| | | *Precision* | *Recall* | *Precision* | *Recall* | |
| | 25 | 1.000 | 1.000 | 0.998 | 0.997 | 39 |
| | 35 | 1.000 | 1.000 | 0.998 | 0.997 | 43 |
| | 50 | 1.000 | 1.000 | 0.998 | 0.997 | 45 |

#### 2. The effect of gap size threshold between simple pairs

Suppose two adjacent simple pairs $s_a = (i_{a,1}, i_{a,2}, j_{a,1}, j_{a,2})$ and $s_b = (i_{b,1}, i_{b,2}, j_{b,1}, j_{b,2})$, we define gaps($S_a$, $S_b$) = $j_{b,1} - j_{a,2}$. If gaps($S_a$, $S_b$) is more than 300bp and the sequences in the gaps are dissimilar, the two simple pairs will be separated into different groups. Here, we investigate the effect of different gap sizes on Sim_Chr1. Table S2 summaries the result. It can be observed that the three different gap sizes performed equally well in terms of variant detection.

*Table S2*. The effect of *gap size threshold* on the Sim_Chr1 dataset.

| ***Sim_Chr1*** | ***Gap size*** | ***SNV*** | | ***Indel*** | | ***Run time (second)*** |
|---|---|---|---|---|---|---|
| | | *Precision* | *Recall* | *Precision* | *Recall* | |
| | 100 | 1.000 | 1.000 | 0.998 | 0.997 | 41 |
| | 300 | 1.000 | 1.000 | 0.998 | 0.997 | 39 |

| | | | | | | |
|---|---|---|---|---|---|---|
| | 500 | 1.000 | 1.000 | 0.998 | 0.997 | 41 |

3. The selected genome comparison tools and their argument setting

Table *S3* lists the argument setting for each method tested in this study.

*Table S3*. Aligner and their arguments used on the benchmark datasets, where fa1 and fa2 are input genomes with FASTA format.

| Genome comparison tool | Arguments |
|---|---|
| GSAlign | GSAlign -i *idx* -q query.fa -t 8 (for benchmark datasets and PanTro4)<br>GSAlign -i *idx* -q query.fa -sen -t 8 (for mouse chromosome 12) |
| Minimap2 | minimap2 -d *idx* ref.fa<br>minimap2 -t 8 -ax asm10 *idx* fa2 > out.sam (for simHG-1x and simHG-3x)<br>minimap2 -t 8 -ax asm20 *idx* fa2 > out.sam (for simHG-5x) |
| MUMmer4 | nucmer --mum --threads=8 --load=*idx* --prefix=out fa1 fa2 (for benchmark datasets)<br>delta-filter -q out.delta > out.filter.delta<br>delta2maf out.filter.delta > out.maf |
| LAST | lastdb -uNEAR -R01 *idx* fa1<br>lastal -P8 -l30 -k20 -E0.01 *idx* fa2 \| last-split –m1 > out.maf |

4. Benchmark datasets and sequence mutation simulator

The sequence mutation simulation (SVsim.cpp), benchmark datasets, and evaluation program (Evaluation.cpp) in this study are available at http://bioapp.iis.sinica.edu.tw/~arith/GSAlign/.

The diploid sequence of NA12878 genome can be downloaded at http://sv.gersteinlab.org/NA12878_diploid/NA12878_diploid_2017_jan7/, and the variants is available at ftp-trace.ncbi.nlm.nih.gov/giab/ftp/release/NA12878_HG001/.
